# Classical statistical methods are powerful for the identification of novel targets for the survival of breast cancer patients

**DOI:** 10.1101/2024.10.24.620147

**Authors:** Benyapa Insawang, Max Ward, Zhaoyu Li, Amittava Datta

## Abstract

Breast cancer is a leading cause of cancer-related deaths among women. The identification of survival-related target genes is critical for improving the prognosis and outcomes of breast cancer patients. Many methods have been applied to this investigation, such as bioinformatics and machine learning approaches, yet few targets identified from these approaches have been applied in clinics. Here, we present a novel approach by using classical statistical methods of Kolmogorov-Smirnov (KS) test and Jensen-Shannon (JS) divergence to analyse the survival time and gene expression data of breast cancer patients (BRCA) from The Cancer Genome Atlas (TCGA). These methods help compare the survival time distributions and differentiate patients into high and low-risk groups based on gene expression profiles. 1,124 survival-related genes were identified based on the KS test and 18 from JS divergence values. We also identified the optimal thresholds of the expression level of these target genes, which enabled the best separation of survival groups for all breast cancer patients and each subtype of breast cancer patients. These targets were further validated through bootstrapping to ensure that significant results are not due to chance. By comparing those survival targets from previous studies, we found two were novel targets, and two were consistent with previous reports. Overall, our study provides a novel approach for identifying survival targets for breast cancer patients by integrating a series of classical statistical methods, such as the KS test, JS divergence, and bootstrapping. Our approach could also be applied to identifying the survival targets for other cancer types and provide valuable insights into cancer research and clinical applications.

## 1 Introduction

Breast cancer’s significant impact is highlighted by its high incidence and mortality rates globally (1). Biomarker identification can be approached using differentially expressed genes (DEGs), machine learning models, or a combination of both (2; 3; 4). However, identifying DEGs in human RNA-seq samples can lead to high false discovery rates when using existing bioinformatics methods, potentially resulting in misleading interpretations (5). Additionally, the need for pre- defined group labels in DEGs analysis and machine learning models may constrain the discovery of novel insights.

Machine learning offers flexibility and power in uncovering complex associations but can create a black box issue where its inner workings remain unclear. While these algorithms focus on predicting outcomes, they don’t establish causal relationships (6). A statistical model is more suitable for this analysis since we aim to extract information to identify biomarkers—without defined group labels.

Research on survival analysis with gene expression has explored categorizing patients into risk groups through methods like median or quantile-based stratification and classical models, including the Cox model, log-rank test, and prognostic index (PI) (7; 2). However, using median or quantile values as dividing points may not capture significant biological distinctions. Relying solely on the median as a cut-off can also lead to losing important information, particularly when data is heavily skewed. Moreover, the Cox model has limitations due to its reliance on the proportional hazards (PH) assumption. This may not hold in cancer progression, where factors like tumour characteristics and treatment responses can vary significantly throughout a patient’s journey (8). To address these limitations, alternative distance metrics could be explored to measure differences in survival times more effectively.

The Kolmogorov-Smirnov (KS) test is a nonparametric method that does not rely on specific distributional assumptions (9). This makes it versatile for comparing the empirical distributions of two samples without assuming a particular distribution form. In addition, the Jensen-Shannon (JS) divergence is another metric used to compare probability distributions. The JS divergence measures the similarity between two probability distributions by calculating the average of the Kullback-Leibler (KL) divergences between each distribution and their average (10). Using these methods, we can determine the gene expression levels, effectively distinguishing patient survival time distributions and overcoming the limitations of median or quantile-based approaches.

This study makes the following contributions:

- We identify biomarkers and expression levels significantly related to the survival of breast cancer patients.
- We introduce a statistical model capable of identifying high- risk genes interpretably, with statistically sound guarantees for correctness.
- We enhance data utilisation and interpretability by using statistical methods, such as the KS test and JS divergence.

## 2 Results

The study flowchart is presented in Figure 1 to identify genes associated with the survival time of breast cancer patients.

**Figure 1.**
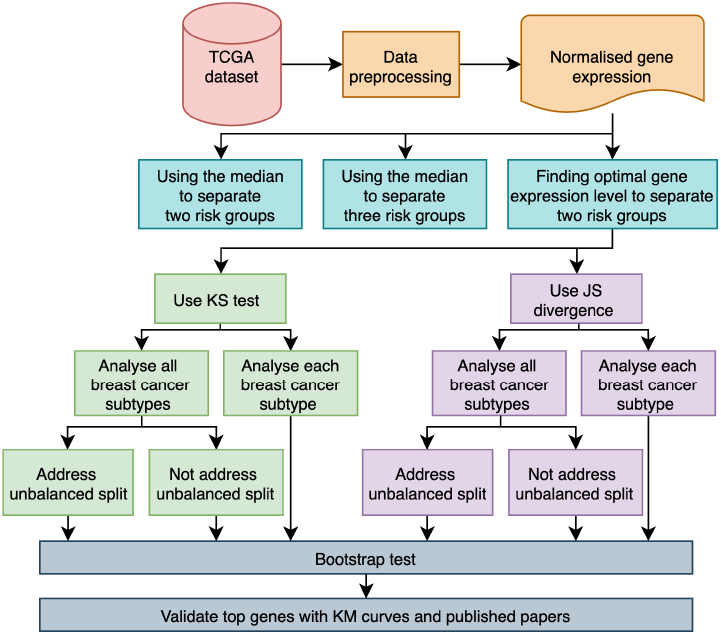
Overview of the study workflow.

### 2.1 Survival-related genes identified by using the KS test

After using statistical methods to find the optimal expression level of each gene, we selected genes with a KS test p-value *≤* 0.01, indicating a significant separation of the two groups’ survival times. We applied bootstrapping to obtain p-values *≤* 0.001 to minimise the risk of false positives. The number of significant genes identified using the KS test is shown in Table 1.

**Table 1.** Number of significant genes based on KS and bootstrapping tests.

| Breast cancer subtype | Number of genes |
| --- | --- |
| All Subtypes | 1,124 |
| LumA | 534 |
| LumB | 288 |
| Her2 | 1,038 |
| Basal | 2,074 |
| Normal-like | 155 |

When we increased the sample size to at least 1%, 10%, 20%, 30%, and 40% in one group, the number of significant genes we found was 257, 93, 38, 16, and 12, respectively. Subsequently, we identified the intersection of these genes to test whether they consistently appear significant regardless of the imbalance, resulting in 12 genes. However, these genes failed the bootstrap test.

Figure 2 demonstrates that certain breast cancer subtypes share significant genes, suggesting underlying common biological mechanisms. Additionally, as shown in Figure 3, there is considerable overlap between the significant genes across all subtypes and those unique to each subtype, highlighting genes that may have roles across multiple breast cancer subtypes.

**Figure 2.**
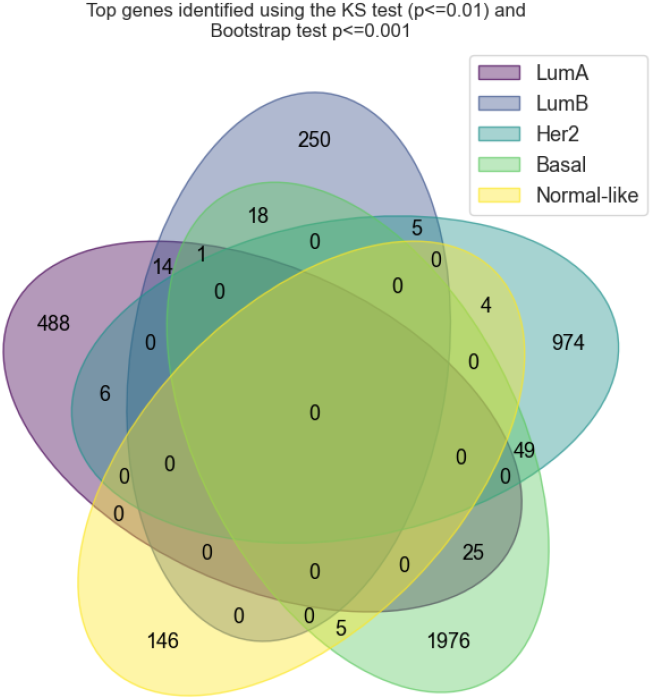
Significant genes identified using the KS test for each breast cancer subtype. Each diagram shows the number of significant genes overlapping with other breast cancer subtypes.

**Figure 3.**
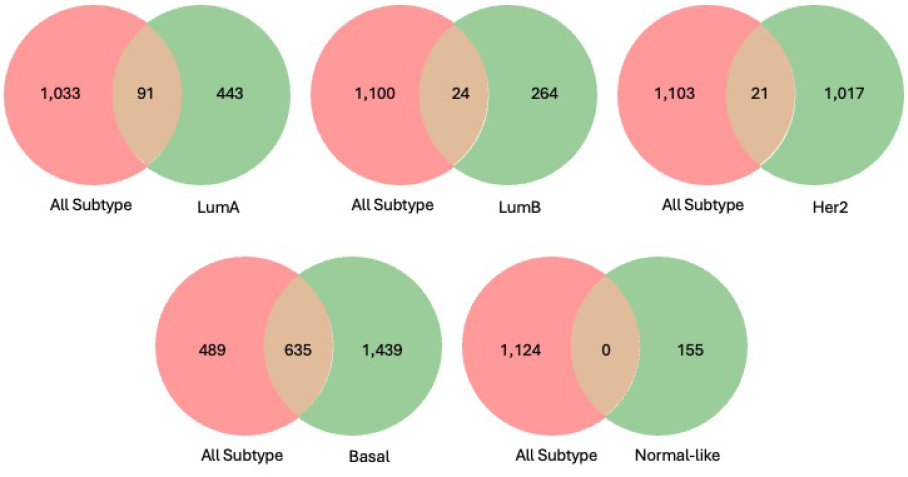
Venn diagram showing the number of significant genes for all breast cancer subtypes overlapping with each subtype from the KS test.

**Figure 4.**
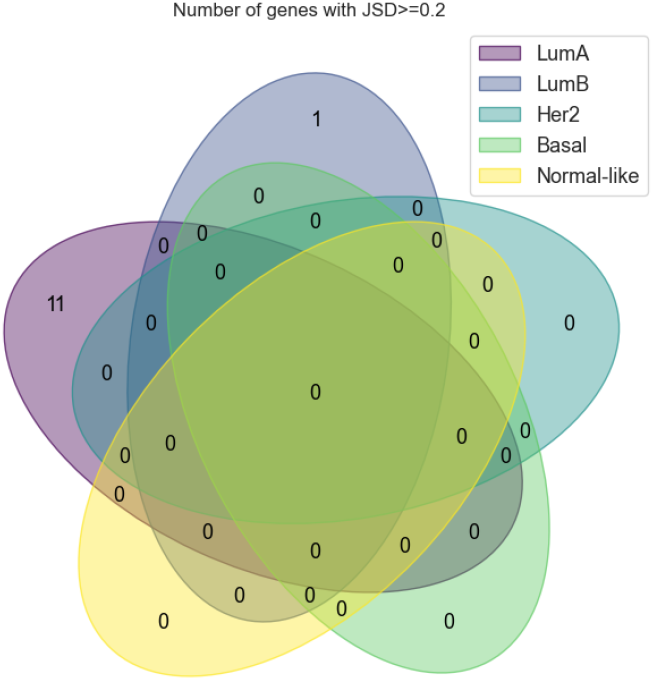
Significant genes identified using JS divergence for each breast cancer subtype. Each diagram shows the number of significant genes overlapping with other breast cancer subtypes.

**Figure 5.**
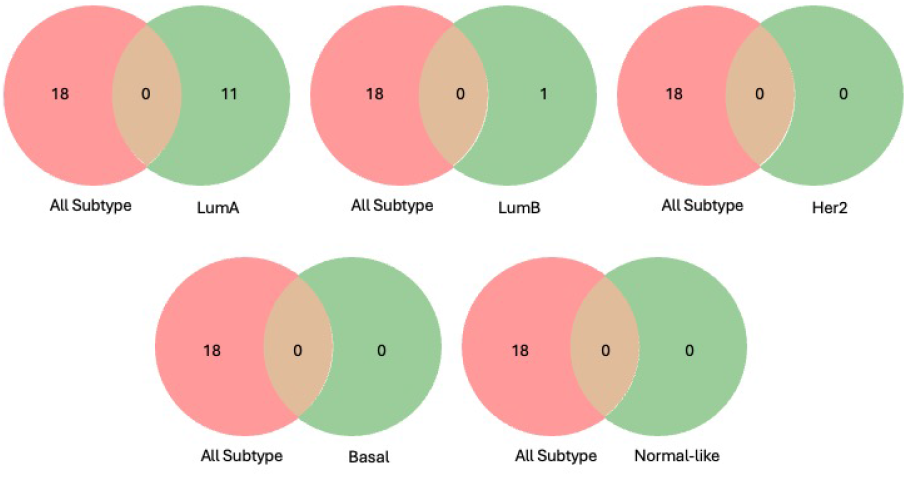
Venn diagram showing the number of significant genes for all breast cancer subtypes overlapping with each subtype from JS divergence.

### 2.2 Survival-related genes identified by using JS divergence

We identified significant genes with a JS divergence of 0.2 or greater and bootstrapped p-values of 0.001 or less. The analysis of JS divergence produced highly unbalanced groupings; therefore, we focused on genes with more balanced survival time distributions. As a result, 18 significant genes were identified across all subtypes: 11 for LumA and 1 for LumB. The Her2, Basal, and Normal-like subtypes yielded no significant genes (Table 2).

**Table 2.**
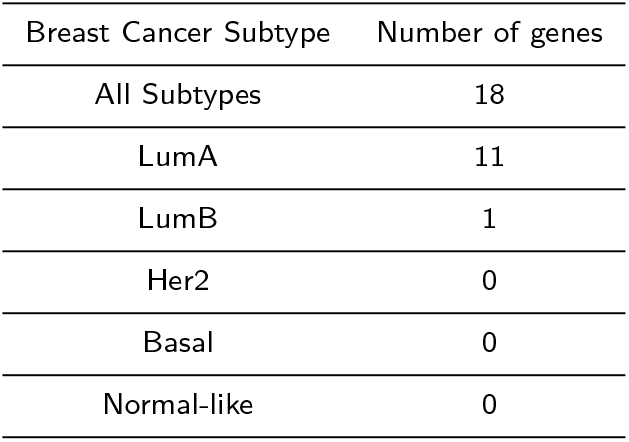
Number of significant genes based on JS divergence and bootstrapping test.

### 2.3 Validation of survival-related genes identified by statistical methods

We found four genes identified by both the KS test and JS divergence: three (DTYMK, LINC01311, PYY2) for all subtypes and one (TMEM222) for LumA (Figure 6). All of these genes are related to cancer. DTYMK is a prognostic biomarker in breast cancer (11). LINC01311 plays a vital role during the tumorigenesis and progression of thyroid cancer (12). PYY2 and TMEM222 have been studied about cancer, but not specifically in breast cancer and survival time. This finding highlights the effectiveness of our methodology as it uncovers published survival-related genes.

**Figure 6.**
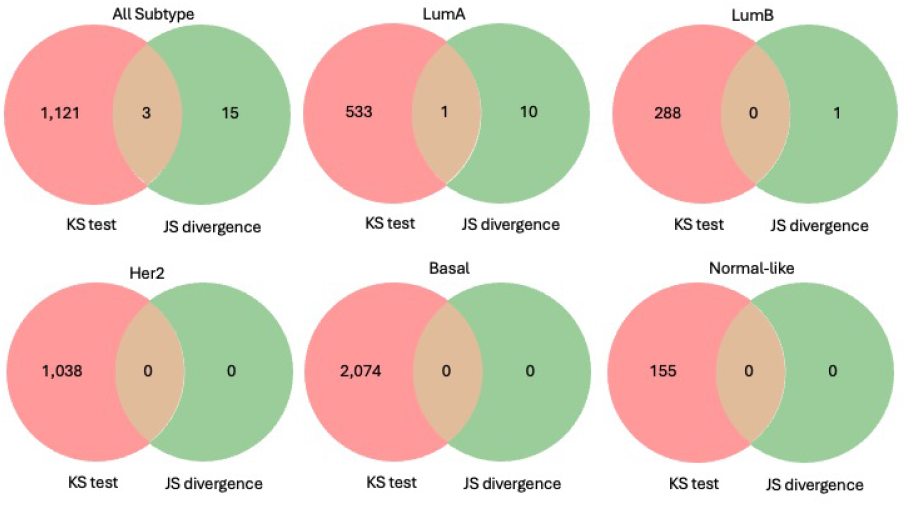
Venn diagram showing the number of significant genes from both KS test and JS divergence.

Figure 7 illustrates the proportion of patients for each breast cancer subtype. The findings align with existing data, showing that patients with LumA and LumB subtypes tend to have better survival rates, which is confirmed by our results.

**Figure 7.**
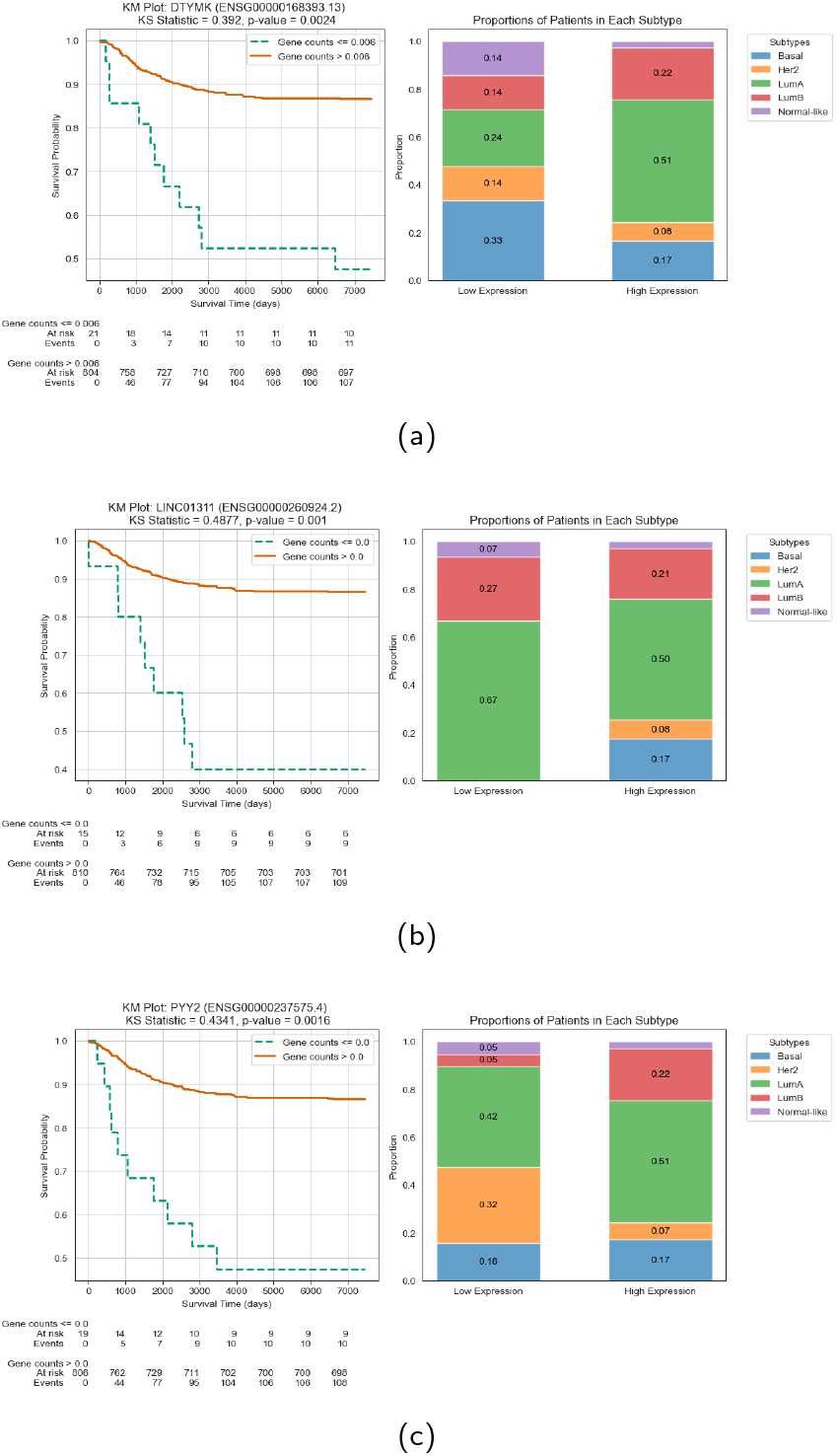
KM curves and proportions of patients in each subtype. (a) KM plot for DTYMK. (b) KM plot for LINC01311. (b) KM plot for PYY2.

### 2.4 Comparison of using the optimal expression level and the median to separate risk groups

We observed improved classification in comparing patient groups using optimal expression levels versus the median. In Figure 8, it is evident that the Kaplan–Meier (KM) curves exhibit improved separation with the optimal cut-off points, demonstrating more distinct survival probabilities between the low- and high-expression groups. This discovery emphasises the significance of employing data-driven methods to identify expression thresholds that enhance the differentiation between risk groups rather than relying solely on the median.

**Figure 8.**
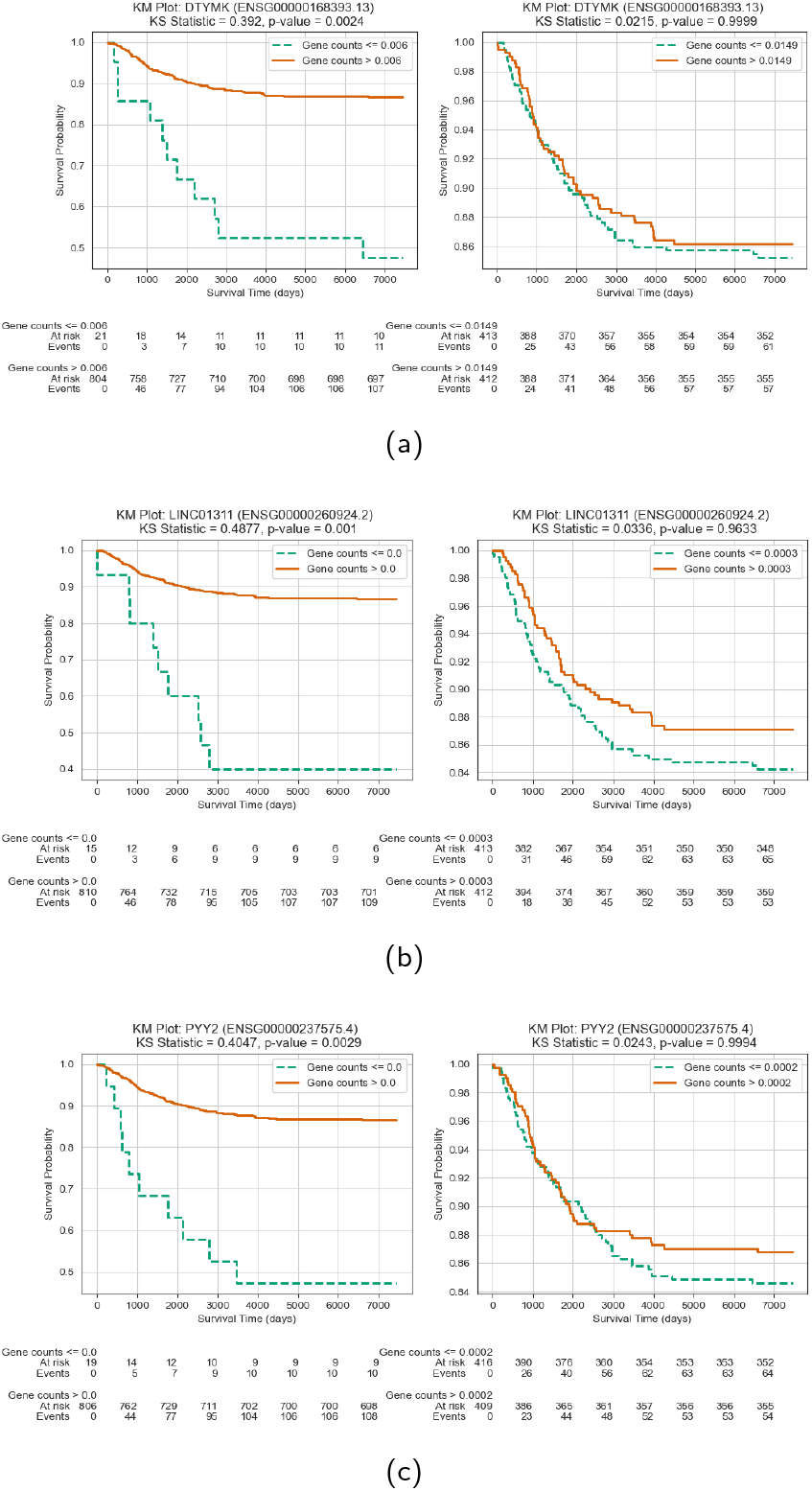
Comparison of KM curves using the optimal expression level (left) and the median (right) to separate risk groups. (a) KM plot for DTYMK. (b) KM plot for LINC01311. (b) KM plot for PYY2.

## 3 Discussion

In our analysis of significant genes using statistical methods, we observed that one of the groups had fewer than 10 patients. This suggests that certain genes may be forming very small groups to achieve a low p-value in the KS test or the highest JS divergence, without considering the number of patients in each group. It is difficult to determine whether this situation actually identifies a significant gene, as establishing the optimal number of patients for each group poses a challenge. To improve our findings, we may need to implement filters or restrictions that ensure low p-values and create more balanced groups.

When we implemented a restriction requiring at least 1%, 10%, or more of the samples in one of the groups, we noticed a decrease in the number of top genes as the groups became more balanced and their distributions became similar. This raises the question: what level of balance is acceptable in this field? To address this, we will need to conduct further experiments.

Using JS divergence to identify optimal gene expression results in a higher rate of false positives. Many significant genes identified by this method have high bootstrap test p-values, indicating that these findings may be unreliable. In contrast, the KS test provides more dependable outcomes, offering consistent results without creating excessively unbalanced group splits. This suggests that the KS test is a more robust method for assessing gene expression differences in our dataset.

Using statistical methods, we can extract reliable insights from small datasets, identifying genes whose expression levels effectively stratify patient risk groups. This approach minimises computational demands and yields results that can be cross-validated with existing studies, enhancing the clinical relevance of the findings. Additionally, robust bootstrapping tests mitigate the risk of false positives, ensuring that the significant genes we identify are not chance occurrences. The survival-related genes identified through this method demonstrate strong correlations with survival time, supported by empirical evidence. This validates our process as a simple yet powerful tool for screening gene lists, offering an alternative to more complex models in future analyses.

## 4 Conclusion

We identified two novel genes, PYY2 and TMEM222, associated with survival rates in breast cancer patients. This underscores the significance of robust statistical methods in uncovering potential biomarkers that could enhance our understanding of cancer biology and patient outcomes.

## 5 Materials and Methods

### 5.1 Data source

The TCGA dataset was obtained from the Genomic Data Commons (GDC) Data Portal. The research cohort was defined by selecting the project as “TCGA-BRCA” for breast cancer and “Experimental Strategy” as RNA-Seq, available at https://portal.gdc.cancer.gov. This study utilised three primary datasets: gene expression counts, clinical information, and metadata regarding breast cancer subtypes.

We used a maximum value of 7,455 days from deceased patients as a censoring point for those who survived. The gene expression values were normalised using four housekeeping genes: GAPDH, ACTB, PUM1, and UBC. A total of 825 patients were studied, with LumA being the most common cancer subtype and having the highest survival rates. The distribution of survival times is significantly skewed to the left, indicating that a larger number of patients experience longer survival while fewer face early mortality.

### 5.2 Using the median to separate risk groups

We divided patients into two risk groups based on the median expression of each gene (*≤* median as low expression and *>* median as high expression). We used the groups to calculate the survival data’s KS statistic and JS divergence. In the KS test, genes were ranked by the smallest p-values, indicating significant differences in survival times. For the JS divergence, genes were sorted by the highest values, reflecting greater dissimilarity in distribution.

### 5.3 Finding optimal gene expression level

We employed the KS test and JS divergence to determine the most effective gene expression level for distinguishing patients into two groups based on lower and higher survival times. We analysed gene expression across all breast cancer subtypes and within each specific subtype. This process resulted in groups with the lowest KS test p-values and the highest values from JS divergence. We resolved an issue where the algorithm created highly unbalanced sample splits by ensuring that the patients in each group made up at least 1%, 10%, 20%, 30%, and 40% of the total patients. Genes with a KS test p-value *≤* 0.01 and JS divergence *≥* 0.2 were reported as significant genes.

### 5.4 Bootstrapping

We conducted a bootstrapping test (13) to assess whether the significant genes were statistically significant under the null hypothesis, which states that there is no difference in the test statistic between the two groups when gene expression is randomised. We calculated one-sided p-values to generate the final gene list for comparison with published studies.

## 6 Author Contributions

Max Ward, Zhaoyu Li, and Amittava Datta conceived and supervised the study. Benyapa Insawang performed the experiments and wrote the manuscript.

